# Phase alignment of low-frequency neural activity to the amplitude envelope of speech reflects evoked responses to acoustic edges, not oscillatory entrainment

**DOI:** 10.1101/2020.04.02.022616

**Authors:** Yulia Oganian, Katsuaki Kojima, Assaf Breska, Chang Cai, Anne Findlay, Edward Chang, Srikantan Nagarajan

## Abstract

The amplitude envelope of speech is crucial for accurate comprehension. Considered a key stage in speech processing, the phase of neural activity in the theta-delta bands (1 - 10 Hz) tracks the phase of the speech amplitude envelope during listening. However, the mechanisms underlying this envelope representation have been heavily debated. A dominant model posits that envelope tracking reflects entrainment of endogenous low-frequency oscillations to the speech envelope. Alternatively, envelope tracking reflects a series of evoked responses to acoustic landmarks within the envelope. It has proven challenging to distinguish these two mechanisms. To address this, we recorded magnetoencephalography while participants listened to natural speech, and compared the neural phase patterns to the predictions of two computational models: An oscillatory entrainment model and a model of evoked responses to peaks in the rate of envelope change. Critically, we also presented speech at slowed rates, where the spectrotemporal predictions of the two models diverge. Our analyses revealed transient theta phase-locking in regular speech, as predicted by both models. However, for slow speech we found transient theta and delta phase-locking, a pattern that was fully compatible with the evoked response model but could not be explained by the oscillatory entrainment model. Furthermore, encoding of acoustic edge magnitudes was invariant to contextual speech rate, demonstrating speech rate normalization of acoustic edge representations. Taken together, our results suggest that neural phase locking to the speech envelope is more likely to reflect discrete representation of transient information rather than oscillatory entrainment.

## Introduction

Speech comprehension is essential to human communication. A major computational step in neural processing of speech is the extraction of its amplitude envelope, the overall intensity of speech across spectral bands. The speech envelope is dominated by fluctuations in the range of ∼1–10Hz, which are temporally correlated with the syllabic structure of speech, and the removal of which from speech severely impairs intelligibility (1,2). Many studies have shown a consistent relationship between the phase of band-limited low-frequency neural activity measured in M/EEG over auditory cortical areas and the phase of the amplitude envelope of speech, a phenomenon widely known as envelope tracking (3,4). The strength of envelope tracking is correlated with speech intelligibility, suggesting that it could constitute an essential stage in speech comprehension (5,6). However, the neural computations underlying speech envelope tracking are controversial (7–9).

A dominant theory of speech envelope tracking posits that it reflects the entrainment (i.e., phase alignment) of endogenous neural oscillations to envelope fluctuations. According to this theory, phase correction is driven by discrete acoustic landmark events in the speech signal, and occurs primarily for oscillators in the delta-theta range (1-10 Hz), matching the syllabic rate of the speech signal (10–12). Functionally, oscillatory entrainment is thought to benefit speech processing via the self-sustaining property of oscillating dynamical systems (13,14), resulting in automatically-driven prediction of the timing of upcoming information (e.g., syllables (15)).

However, recent work has demonstrated that phase alignment of low-frequency neural activity can be the outcome of transient neural responses rather than oscillatory dynamics (16,17). This becomes pertinent in the case of speech, as it has been suggested that the speech envelope is encoded in evoked responses to the same acoustic landmark events that supposedly drive the entrainment process. Recent electrophysiology recordings suggest that these events are peaks in the rate of amplitude envelope change that mark the perceived onset of vowels. Yet to date, the phase adjustments predicted by this non-oscillatory process, and whether speech envelope tracking is better explained by it or by an oscillator-based process, remain unclear. The two competing models have drastically disparate functional and mechanistic implications (18–21).

To address this, we combined a model-based computational approach with neurophysiological (MEG) recordings of neural responses in an ecologically valid context, using natural continuous speech. We implemented an oscillatory entrainment model and an evoked responses model, quantified the spectral content and temporal dynamics of neural activity predicted by each model in response to speech, identified diverging model predictions, and tested them against MEG data.

Our modelling approach had two critical features. First, we analyzed phase patterns as event-locked, relative to acoustic landmarks events that are thought to drive both entrainment and evoked response mechanisms. This allowed us to have an extremely high number of events (2106 within participant), and to probe phase alignment in a time-resolved manner. Particularly, it enabled us to quantify reverberation following a phase-reset, a hallmark of oscillatory processes. Second, we additionally presented continuous speech at, equally intelligible, 1/3 of its original rate. In natural speech, the speech rate, and hence the expected frequency of an entrained oscillator, overlaps with the spectral content of evoked responses. Moreover, the duration of an evoked response is longer than the time between phase-resetting events, where oscillatory reverberation is expected to occur. We hypothesized that slowing speech would separate the frequencies of phase alignment between the two models and would space phase-resetting events to distinguish between phase alignment due to evoked responses and extended reverberation.

Comparing cortical dynamics for regular and slowed speech also allowed us to address the neural mechanisms of speech rate normalization. Speech rate normalization allows listeners to adjust perceptual processes to differences in speech rate. It has previously been proposed that speech rate normalization relies on shifts in the frequency of the phase-locked oscillator to the speech rate (3,22–25). Here we examined this hypothesis in naturalistic speech.

## Methods

### Participants

Twelve healthy, right-handed volunteers (six females; age range 22-44 years, median 25 years) participated in the study. All participants were native speakers of English. All participants provided informed written consent and received monetary compensation for their participation. The study was approved by the University of California, San Francisco Committee on Human Research.

### Speech stimulus

Participants listened to two stories (one male, one female speaker) from the Boston University Radio Speech Corpus (BURSC, Table S1) (26), each once at regular speech rate and once slowed to 1/3 speech rate. Overall, the stimuli contained 26 paragraphs (each containing 1 - 4 sentences) of 10 – 60 s duration, with silent periods of 500 – 1100 ms inserted between paragraphs to allow measuring onset responses in the MEG without distortion from preceding speech. Boundaries between paragraphs corresponded to breaks between phrases, such that silences were perceived as natural. Speech stimuli were slowed using the Pitch Synchronous Overlap and Add (PSOLA) algorithm, as implemented in the software Praat (27), which slows down the temporal structure of the speech signal while keeping its spectral structure constant (28). Overall, the regular speech stimulus was 6.5 min long and the slowed stimulus was 19.5 min long.

### Procedure and stimulus presentation

All stimuli were presented binaurally at a comfortable ambient loudness (∼ 70 dB) through MEG compatible headphones using custom-written MATLAB R2012b scripts (Mathworks, https://www.mathworks.com). Speech stimuli were sampled at 16 kHz. Participants were asked to listen to the stimuli attentively and to keep their eyes closed during stimulus presentation.

Participants listened to the radio stories once at regular and once at slowed rate in separate but interleaved blocks, such that each participant heard one story first at regular speech rate and the other at slowed speech rate. Comprehension was assessed with 3-4 multiple choice comprehension questions posed after each story (Table S2). For each participant, a different randomly selected subset of questions was used for each block. Percentage correct was compared between regular and slow blocks using a two-sided paired t-test.

### Neural data acquisition and preprocessing

MEG recordings were obtained with a 275-axial gradiometers whole-head MEG system (CTF, Coquitlam, British Columbia, Canada) at a sampling rate of 1,200 Hz. Three fiducial coils were placed on the nasion and left and right pre-auricular points to triangulate the position of the head relative to the MEG sensor array. The position of the patient’s head in the device relative to the MEG sensors was determined using indicator coils before and after each recording interval to verify an adequate sampling of the entire field. The fiducial markers were later co-registered onto a structural magnetic resonance imaging scan to generate head shape (29).

### Data analysis and modelling

All analyses were conducted in MATLAB R2019a (Mathworks, https://www.mathworks.com) using custom-written scripts and the FieldTrip toolbox (30).

### Acoustic feature extraction

We extracted the broad amplitude envelope of speech stimuli by rectifying, low-pass filtering at 10 Hz and down-sampling to 100 Hz of the original stimulus waveform in this order. We then calculated the derivative of the resulting loudness contours as a measure of the rate of change in the amplitude envelope. Finally, we extracted the sparse time series of local peaks in the amplitude envelope (peakEnv) and its derivative (peakRate). All features are depicted in Figure 1A, for an example stimulus excerpt. Overall, the stimulus set contained 2106 peakRate and 2106 peakEnv events per speech rate condition.

**Figure 1.**
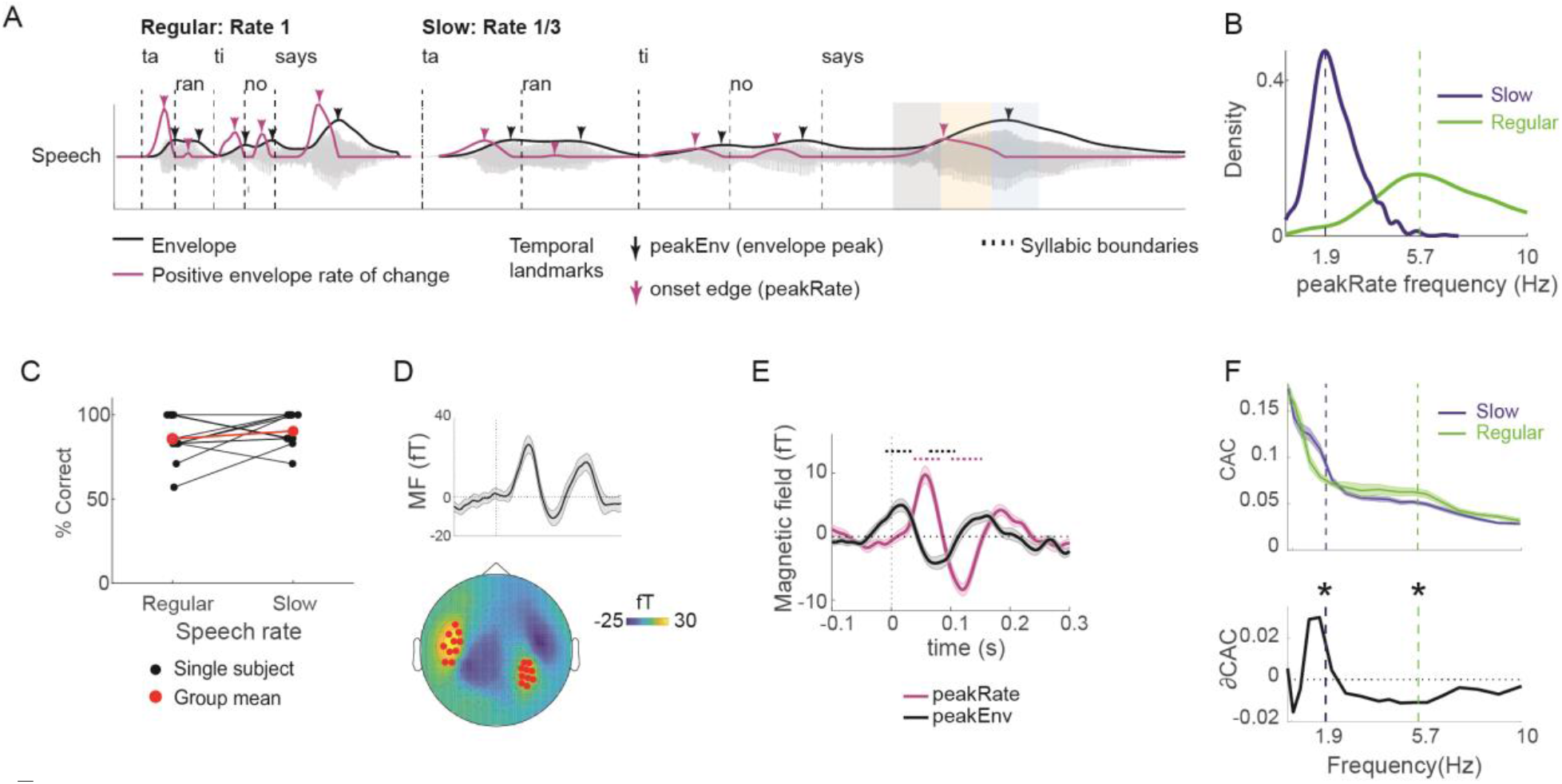
Task design and envelope tracking in neural data. **A**. The acoustic waveform of an example utterance (‘Tarantino says…”), with syllable boundaries, amplitude envelope, and first temporal derivative of the envelope superimposed on it. The same utterance is shown at a regular rate (left) and slowed (right) speech rate. Arrows mark candidate temporal landmark that might induce phase locking (Black: local peaks in the envelope, peakEnv; Purple: acoustic edges, defined as local peaks in the first temporal derivative (rate of change) of the envelope, peakRate). **B**. Frequency of occurrence for peakRate/peakEnv events. Dashed vertical lines mark the average frequency of peakRate events in slow (blue, 1.9 Hz) and regular speech (green, 5.7 Hz). **C**. Single-subject (black) and group-average (red) comprehension performance. **D**. Sensor selection was based on M100 response to utterance onsets. Top: Group-averaged evoked response across all 20 sensors included in the analysis. Error bars are ± 1 SEM across subjects. Bottom: Topographic map of a group-averaged M100 response with selected sensors marked in red. **E**. Group-averaged evoked response aligned to peakRate and peakEnv events. Dotted lines mark clusters with *p* < 0.05 with a cluster-based permutation test against 0. Error bars are ± 1 SEM across subjects. **F**. Cerebro-acoustic phase coherence (CAC) between MEG responses and speech envelope (upper panel), and the difference between slow and regular speech (ΔCAC, lower panel). Data were filtered in semi-logarithmically spaced bands between 0.3 and 10 Hz for this analysis. Dashed vertical lines mark the average frequency of peakRate events in each condition, as shown in D. * *p* < 0.01 in post-hoc t-tests with interaction *p* < 0.01. Error bars are ± 1 SEM across subjects.

### Evoked response and oscillatory entrainment models for IEPC simulation

We implemented two computational models that predict neural activity in response to continuous speech, one based on oscillatory entrainment and another based on evoked responses. We then submitted their output to the same phase analysis as for MEG data. We assumed that both processes were driven by peakRate events, based on our analysis of responses to acoustic landmarks and previous work (Oganian & Chang, 2019). As input, each model received a time series that contained peakRate values, scaled within speech rate to between 0.5 and 1, at times of peakRate events, and zeros otherwise. We scaled to this range as our analyses revealed that neural phase alignment to speech is normalized within each speech rate, and that its magnitude for the bottom quantile is ∼50% of the top quantile (see Results, Figure 5). To capture the variable latency of the neural response to non-transient sensory events such as acoustic landmarks, we added random temporal jitter (gaussian distribution, SD = 10 and 30 ms in regular and slow speech, respectively) to the timestamp of each peakRate event. Subsequent phase analyses were conducted using the original, non-jittered time stamps. To account for the non-uniform spectral impact of the 1/f noise that is typical to neurophysiological measurement, we added noise with this spectral content to the predicted neural response output by each model, with a signal-to-noise ratio of 1/10. To create the noise, we filtered gaussian white noise to the 1/f shape with the Matlab function firls.m. The temporal and amplitude jitter parameters were fitted to maximize the similarity between the predicted and observed spectrotemporal patterns of phase alignment. Importantly, to not favor one model this was done across the two models and speech rates. To ensure that results would not be biased by the introduction of simulated random noise, we repeated the randomization procedure 2560 times for each model and each speech rate (64 iterations of temporal noise X 40 iterations of amplitude noise), calculated the phase analyses (below) on the predicted neural signal from each randomization, and then averaged across randomizations.

For the oscillator model, peakRate events induce phase correction of a fixed-frequency oscillator whose frequency is centered on the speech rate (5.7 and 1.9 Hz for regular and slow speech, respectively), as is assumed by oscillatory entrainment models and confirmed in previous work (16,31). Following Large & Snyder (31), this process was modelled using a coupled oscillator dynamical system:

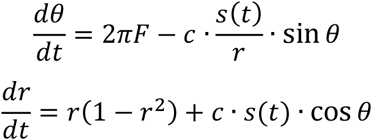

The system produces periodic limit cycle behavior at a radius of *r* = 1 (attractor point) and a frequency *F* in the absence of input (*s(t)* = 0) and follows phase correction towards an angle of θ = 0 when presented with input (*s(t)* > 0). The magnitude of phase correction depends on the strength of the input, the current angle, and the coupling parameter *c*. At low values of c, no oscillator was able to entrain to speech, whereas at high values, entrainment spread across all oscillator frequencies. Crucially, as predicted, at intermediate values, only the oscillator with the correct frequency was entraining to our speech stimulus (Fig. 2B). We thus focused on an oscillator model with intermediate entrainment strength and oscillator frequency corresponding to the speech rate in each task condition for further analyses. Specifically, the value of *c* was set such that the maximal phase correction possible (when *s(t)* = 1 and 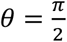or 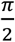 would be 70% of the maximal phase shift. We reconstructed the predicted response as: *PredRespi* = *cosθ*_*i*_ *· r*_*i*_.

**Figure 2.**
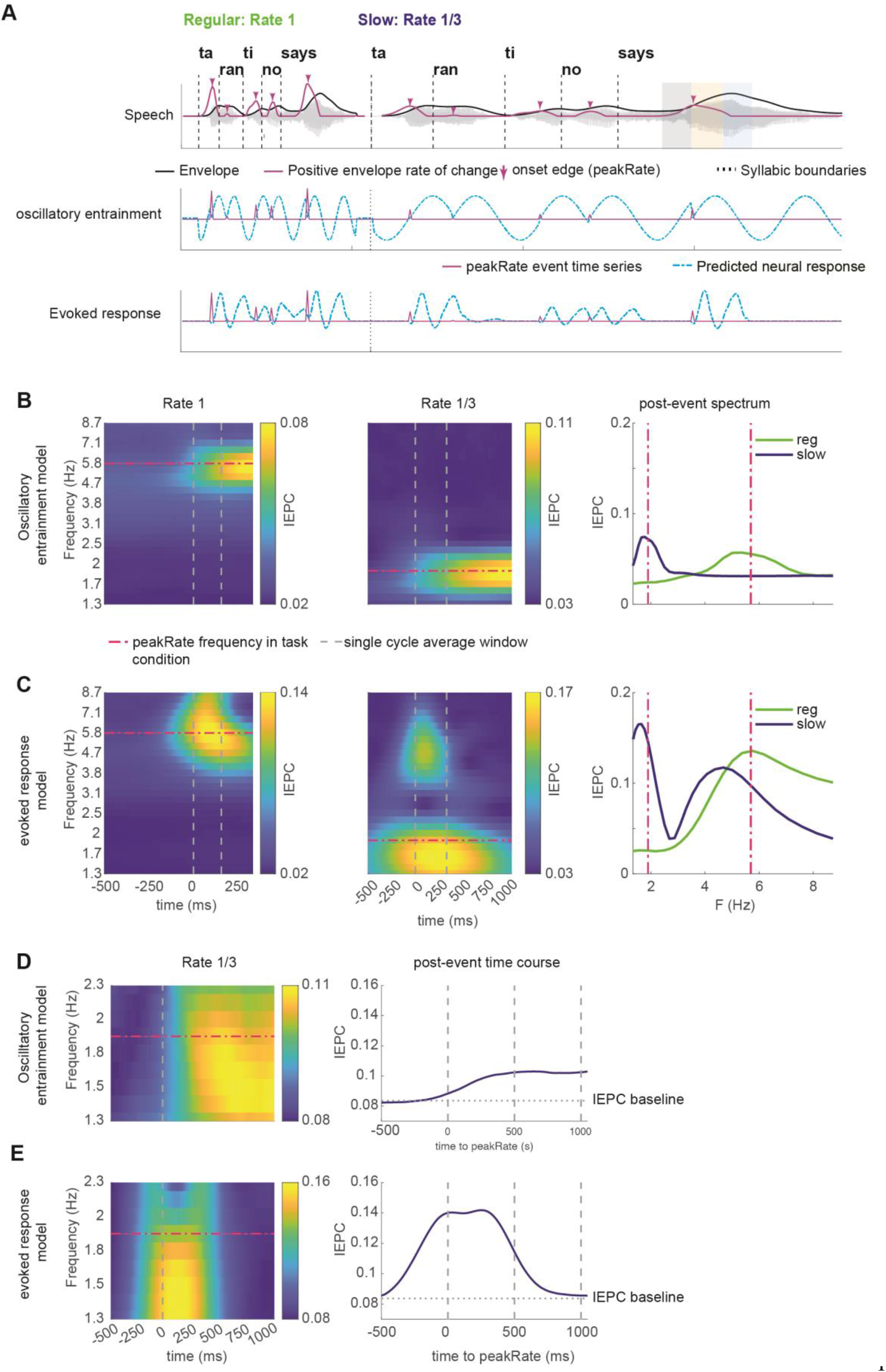
Spectral and temporal signatures of inter-event phase coherence (IEPC) in oscillatory entrainment and evoked response models. **A**. Schematic illustrations of the predicted neural response to the utterance in Figure 1A using three different models. Top: speech signal. Middle: oscillatory entrainment model; Bottom: Evoked response model. **B**. IEPC patterns predicted by oscillatory entrainment model for regular and slow speech with a focus on spectral precision. Dashed lines indicate the frequency of peakRate events in each condition. **C**. As B for evoked response model. **D**. Temporal dynamics of delta-IEPC predicted by oscillatory entrainment model, based on peakRate events that are at least 1000ms apart from following events (n = 113 events in slow speech). **E**. Same as D for slowed speech.

For the evoked response model, peakRate events trigger a prototypical evoked response with its amplitude proportional to the strength of the input. This process was modelled using a linear convolution of the time series of peakRate events with the waveform of an evoked response to peakRate events. The latter was estimated directly from the MEG data, using a time-delayed linear encoding model (Temporal Receptive Field, TRF (32,33)), with a time window of −150 to 450 ms relative to peakRate events. While we found no effect of speech slowing on the shape of the neural response to peakRate events in our previous intracranial work (32), we assumed that neural responses recorded with MEG will be additionally shaped by other speech features that occur in temporal proximity to peakRate events (e.g., vowel onsets). Therefore, we estimated the evoked response separately within each speech rate. We used the TRF approach instead of simple averaging due to the high rate of peakRate events (average interval ∼170 ms), which would have distorted the averaging-based estimate due to overlap between evoked responses.

### MEG data preprocessing

Offline data preprocessing included (in this order) artifact rejection with dual signal subspace projection (DSSP) and down-sampling to 400 Hz. DSSP is a MEG interference rejection algorithm based on spatial and temporal subspace definition (34). Its performance has been recently validated using clinical data (35). In all subsequent analyses of segmented data, segments containing single sensor data above 1.5pT and visually identified artifacts (including muscle, eye blink, and motion) were flagged as bad events and removed from further processing (0.2 % of segments).

### Event related analysis and sensor selection

For all evoked response analysis, we first extracted the broadband signal by band-pass filtering the data between 1 and 40 Hz (second-order Butterworth filter). To focus all further analyses on responses originating in temporal auditory areas, we selected sensors based on the magnitude of the group averaged M100 response to the onset of utterances (independent of responses to acoustic features within the utterance, which were the focus of subsequent analyses). For this purpose, we segmented the broadband signal around utterance onsets (−200 to 500 ms), averaged these epochs across utterances and participants, applied baseline correction (−200 ms to 0 ms relative to utterance onset), and extracted the M100 amplitude as the average activity between 60-100 ms after utterance onset. We then selected the ten sensors with maximal M100 responses from each hemisphere, and all subsequent analyses were conducted on these 20 sensors.

To identify which landmark in the speech envelope drives evoked responses, we analyzed evoked responses to peakRate and peakEnv events. We reasoned that with alignment to an incorrect landmark, evoked responses would have reduced magnitude due to smearing, and latency that is shifted away from the acoustic event. For this purpose, we segmented the broadband signal around acoustic landmark events (−100 to 300 ms), averaged these epochs across events within each participant separately for peakRate and peakEnv events, and applied baseline correction (−100 ms to 0 ms relative to event onset). Based on our previous work (REF), we hypothesized that peakRate events would be the driving acoustic landmark. We compared evoked responses to peakRate and peakEnv using timepoint by timepoint t-tests.

### Time-Frequency decomposition

Identical time-frequency (TF) analyses were performed on the continuous MEG data and on the continuous simulated signal from the Evoked Response and Oscillatory Entrainment models. To evaluate the instantaneous phase of the signal at individual frequency bands (logarithmically spaced between 0.67 and 9 Hz, 0.1 octave steps), we applied non-causal band-pass Butterworth filters around each frequency of interest, performed the Hilbert transform, and obtained the amplitude and phase as the absolute value and phase angle, respectively, of the Hilbert signal. Filter order was chosen to achieve maximal 3 dB of passband ripple and at least 24 dB of stopband attenuation. We conducted this TF analysis with a narrow filter width (±0.1 octave of the frequency of interest) for analyses of spectral patterns to increase frequency resolution, and again with a wider filter (±0.5 octave) for analyses of temporal dynamics to increase temporal resolution.

### Cerebro-acoustic phase coherence (CAC)

To assess cerebro-acoustic phase coherence between the speech envelope and MEG responses, the speech envelope was processed using the same procedure that was applied to the MEG responses: Down-sampling and TF analysis using the wide filter settings. Phase locking between the speech envelope and MEG response was calculated across the entire duration of every utterance within each frequency band, using the Cerebro-acoustic phase coherence (CAC):

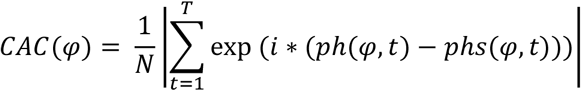

where φ is the center frequency of a frequency band, *T* is the number of time samples in an utterance, *ph* is the phase of the neural signal, and *phs* is the phase of the speech envelope in band φ at time *t*. To equate the number of time points entering the analysis for slow and regular speech, slow speech utterances were split into three equal parts before CAC calculation, and resultant CAC values were averaged. CAC was averaged across sensors for each hemisphere.

A priori, we hypothesized that CAC would differ between conditions in the frequency bands corresponding to the average frequency of peakRate events in each rate condition (regular: 5.7 Hz; slow: 1.9 Hz, Figure 2D). We tested this hypothesis using a 3-way repeated-measures ANOVA with factors frequency band (high/low), factor speech rate (slow/regular), and hemisphere (left/right). To test for further differences in each frequency band, we assessed the effect of speech rate and hemisphere onto CAC using a two-way repeated-measures ANOVA with factor speech rate (slow/regular) and hemisphere (left/right). Significance in this analysis was Bonferroni-corrected for multiple comparisons across bands.

### Inter-event phase coherence (IEPC)

Both IEPC analyses were conducted on the actual MEG data and the neural responses predicted by the evoked response and oscillatory entrainment models. To assess neural phase locking around peakRate events, we segmented the continuous phase data around peakRate events (see below), and obtained a time-resolved inter-event phase coherence (IEPC) (Lachaux, Rodriguez, Martinerie, & Varela, 1999). For each timepoint, IEPC was calculated using the following formula:

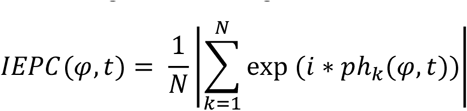

where *N* is the number of events, *ph* is the phase of the neural signal in trial *k*, for the frequency band φ and timepoint *t*. IEPC were first calculated within each of the selected sensors, then averaged across sensors.

#### Spectral patterns of IEPC

To assess the spectral distribution of phase-locking following peakRate events with increased frequency resolution, we segmented the phase data outputted by the narrow filter TF analysis around peakRate events (−500 to 500 ms) and calculated the IEPC. To prevent distortion of the estimated phase by subsequent peakRate events, we only used ones that were not followed by another peakRate event within the 0-500 ms window (n=813 within each participant). To identify whether in this time window and frequency range there was a significant increase in IEPC in the MEG data, the resulting time x frequency IEPC was compared with the pre-event baseline using 2-D cluster-based permutation t-tests (36) with 3000 permutations, a peak t threshold of *p* < 0.01, and a cluster threshold of *p* < 0.01. Baseline IEPC was calculated as the average IEPC between −400 ms to −100 ms relative to event onset in each frequency band.

To compare between model predictions and data, IEPC spectral profiles were calculated, separately for each speech rate condition, by averaging IEPC TF images following peakRate event onset across a time window that conforms to one cycle of an oscillator whose frequency matches the speech rate, i.e. 0-170 ms at regular speech rate and 0 – 500 ms at slowed speech rate.

#### Temporal extent of IEPC

To assess the temporal extent of IEPC between peakRate events, we focused on the slowed speech condition, where phase-locking originating from the evoked response and from putative oscillatory entrainment occupy distinct spectral bands. We segmented the phase data outputted by the broad filter TF analysis around peakRate events (−500 to 1000 ms). with a temporal interval of more than two oscillatory cycles for half an octave around the frequency of peakRate events (1.9 Hz) - that is at least 1040 ms to the next peakRate (n = 114 peakRate events per participant). As this analysis was focused on the temporal dynamics of IEPC, we examined IEPC dynamics as a function of time, averaged across single frequency bands in this range. For the MEG data, this time course was tested against a theoretical chance level, defined as the expected IEPC value for randomly sampling a matched number of angles from a uniform Von-Miese distribution.

### Effect of peakRate magnitude on IEPC

In each rate condition, peakRate events were split into five quantiles, and IEPC was separately calculated within each quantile. Then, we extracted the average IEPC in the theta band (4 – 8 Hz) across all the time points for one cycle of the given frequency band after the event. IEPC in each quantile was compared using 2-way ANOVA with factors quantile and speech rate (regular speech, slow speech).

## Results

### Speech Envelope Tracking for regular and slow speech as seen in MEG

We recorded MEG while participants (*n* = 12) listened to continuous speech containing 2106 envelope landmarks, once at the original rate (Regular speech condition 6.5 minutes duration), and once slowed to 1/3 of the original speech rate (Slow speech condition, 19.5 minutes duration, Figure 1A). Stimulus materials were selected from the Boston University Radio Speech Corpus, BURSC (Ostendorf, Price, & Shattuck-Hufnagel, 1995); see Tables S1 and S2 for full stimulus transcript and list of comprehension questions. Stimuli were split into 26 utterances of 10-69 seconds duration (30 – 210 s in Slow speech condition), with additional silence periods inserted between them. This allowed us to estimate an auditory evoked response to speech onset from the data, without altering the original temporal dynamics of the stimulus within sentences.

In a first step, we characterized the temporal dynamics of acoustic landmark events in our speech stimulus (Figure S1), focusing on peaks in the rate of envelope change (peakRate, n = 2106 per condition, Figure 1A) and on peaks in the envelope (peakEnv, n = 2106 per condition, black in Figure 1A). In the regular speech condition, the average frequency of landmarks (similar for peakRate and peakEnv) was 5.7 Hz (SD = 2.9 Hz, Figure 1B), as is typical in natural speech (Ding et al., 2017). In the slow speech condition, the average frequency of landmarks was 1.9 Hz (SD = 1 Hz, similar for peakRate and peakEnv), shifting the peak of the envelope power spectrum to the delta band. Slowing did not impair participants’ comprehension, as probed by multiple choice comprehension questions after each story (3-4 questions per story, chance-level per question: 50 %; accuracy in regular speech: mean = 83%, SD = 13%; accuracy in slow speech: mean = 90%, SD = 9.5%; *t*(11) = −1.85, *p* = 0.09; Figure 1C).

### Acoustic edges drive MEG evoked responses

We first asked which landmark in the speech envelope drives evoked responses and phase locking to the envelope in regular speech. To focus our analyses on sensors that capture auditory sensory processing, we selected ten sensors with the largest M100 response to speech onsets after silence periods from each hemisphere for all further analyses (Figure 1D). The M100 response showed the typical dipole pattern in each hemisphere (37). First, we examined the characteristics of evoked responses (band-pass filtered 1-40 Hz and averaged in the time domain) locked to peakRate and peakEnv landmark events. While peakEnv closely follows on peakRate in regular speech, the interval between them varies. Thus, aligning to the incorrect landmark should lead to (1) a reduced magnitude of the averaged evoked neural signal due to smearing, and (2) shifts in response onset times away from the acoustic event. We found transient evoked responses with both alignments (Figure 1E). Crucially, the evoked response was of larger magnitude with aligned to peakRate than to peakEnv (peak magnitude: *t*(11) = 5.9, *p* < 0.001). Moreover, this response started after peakRate events, but before peakEnv events (response latency relative to the event for peakEnv: −12.5 ms; peakRate: +50 ms, determined as the first significant time point in a cluster-based permutation test against 0). Together, these analyses indicated that peakRate events, that is, acoustic edges, rather than peakEnv events, that is, envelope peaks, triggered the evoked response in MEG, in line with previous results (32,38,39).

### Cerebro-acoustic phase coherence between speech envelope and MEG

To confirm that cortical speech envelope tracking was present in our data (40), we calculated the cerebro-acoustic phase coherence (CAC) between neural responses and the speech envelope in frequency bands below 10 Hz. CAC is typically increased at the frequency corresponding to the speech rate (25), which in our data corresponds to the frequency of peakRate in each rate condition (regular: 5.7 Hz, slow: 1.9 Hz). Indeed, speech rate had opposite effects on CAC in these two frequency bands (repeated-measures ANOVA, interaction F(1, 11) = 31.20, p < 0.001, *n*^2^ = 0.30, Figure 1F). At 5.7 Hz, CAC was higher for regular speech (*t*(11) = 5.6, *p* < 0.001, *n*^2^ = 0.42), while at 1.9 Hz it was higher for slow speech (*t*(11) = 3.4, *p* = 0.006, *n*^2^ = 0.29). Moreover, CAC was overall higher at lower frequencies (*F*(1, 11) = 16.44, *p* < 0.001, *n*^2^ = 0.39), as is typical for this measure (REF). No other frequency band showed a significant effect of speech rate on CAC (all Bonferroni-corrected p > 0.05). Overall, this result replicates previous findings of cortical speech envelope tracking in frequency bands corresponding to the speech rate of the stimulus. However, as this measure is calculated across the entire stimulus time course, it cannot capture local temporal dynamics in neural phase, driven by phase resets at acoustic edges. To evaluate local temporal and spectral patterns of neural phase-locking following peakRate events, we calculated inter-event phase coherence (IEPC) across peakRate events in the speech stimulus. In contrast to prior studies of CAC, which quantified phase consistency across time, IEPC is calculated across single event occurrences (i.e., single trials) for each time point. IEPC thus enables tracking of the temporal dynamics of phase locking (Gross et al., 2013).

### Oscillator and evoked response models predict distinct patterns of phase alignment to slowed natural speech

To obtain a quantitative estimate of neural phase patterns predicted by oscillatory entrainment and evoked response mechanisms, we implemented computational models of neural envelope tracking as predicted by both processes (see methods for a full description of both models). The input to both models was the acoustic stimulus reduced to peakRate events: a continuous time-series down-sampled to match the MEG sampling frequency and containing non-zero values corresponding to peakRate magnitudes at times of peakRate events, and 0 otherwise. The oscillator model was implemented as a coupled oscillator dynamical system with a non-decaying amplitude attractor point, that followed phase resetting whenever the input was different from 0 (at peakRate events), at a magnitude determined by an entrainment parameter (Breska & Deouell, 2017). A preliminary analysis verified that indeed an oscillator whose endogenous frequency corresponds to the average rate of the speech stimulus would be best suited to entrain to it. The evoked response model was designed as a linear convolution of the peakRate event time series with a stereotypical evoked response, which was extracted from the actual MEG data using a time-lagged linear encoding model (rather than simulated to have an ideal shape) (32,33). To both models, we added 1/f shaped noise, as is observed in neurophysiological data, and a temporal jitter around peakRate event occurrence to each model. See methods for a full description of both models. Both models output a predicted neural response time series (Fig. 2A), from which we extracted predicted spectral and temporal patterns of inter-event phase coherence (IEPC) in the theta-delta frequency ranges following peakRate events for each condition (Fig. 2B).

To identify distinct predictions of the two models, we focused on two aspects of the overall predicted pattern of IEPC. First, we quantified the spectral shape of predicted responses, by examining the average IEPC pattern in the first oscillatory cycle after peakRate events. We found that in regular speech, both the evoked response model and the oscillatory model predicted a transient increase in theta IEPC following peakRate events (Figure 2C+D, left). However, their predictions for the slow speech condition diverged significantly (Figure 2C+D, middle). The oscillator model predicted a single peak in IEPC around the oscillator frequency in IEPC (Figure 2C, right). In contract, the evoked response model predicted two IEPC peaks, around 5.7 Hz and around 1.9 Hz, reflective of the shape of the evoked response and its frequency of occurrence (i.e., the frequency of peakRate events), respectively (Figure 2D, right). We verified this by manually morphing the shape of the evoked response and the frequency of evoked responses, which shifted the location of the upper and lower IEPC peaks, respectively.

Second, we examined the temporal extent of IEPC predicted by each model. A key feature of an oscillatory entrainment mechanism, that is central to the cognitive functions ascribed to oscillatory models, is that the endogenous oscillator will continue to reverberate after phase reset beyond the duration of a single oscillatory cycle, resulting in increased phase alignment for a prolonged time window (13,14,41). In our data, this should be expressed as an increase in IEPC extending beyond a single oscillatory cycle after peakRate events. In contrast, if phase locking is the result of evoked responses to peakRate events, the increase in IEPC should be limited to the duration of an evoked response. To quantify this, we focused our analysis on the first two cycles after peakRate events. To prevent interference from subsequent phase-resetting events, we only included peakRate events that were not followed by another peakRate event in this interval (n=114). Importantly, such events were distributed throughout the speech stimulus and not limited to sentence or phrase ends. As in regular speech rate the duration of the evoked response (∼350 ms, Figure 1D) extends across two putative cycles at the speech rate frequency (∼350 ms at 5.7 Hz), which would not allow to dissociate the two models, we focused this analysis on the slow speech condition. We then examined the time course of IEPC in a range of frequencies surrounding 1.9 Hz, the frequency of the putative oscillator that best entrains to the slow speech rate. As expected, we found divergent predictions: the oscillator model predicts that IEPC remains increased for multiple oscillatory cycles (Figure 2D). In contrast, the evoked response model predicts that the increase in IEPC is temporally limited to the duration of a single evoked response (Figure 2E). Taken together, this model comparison identified two divergent predictions for IEPC patterns in slow speech: The spectral distribution of IEPC and its temporal extent. Next, we performed these identical analyses on our neural data and compared the patterns in the data with the models’ predictions.

### Spectral pattern of Delta-Theta phase-locking to acoustic edges is best described by the evoked response model

We next turned to testing the two divergent predictions of the two models against MEG data, starting with predictions for spectral distribution. Based on the models’ predictions (Fig. 2 and Fig. 3A), we first took a hypothesis-based approach, analyzing average IEPC in predefined time-frequency ROIs: Within a single oscillatory cycle post peakRate event in the theta (4-8 Hz) and delta (1 – 3Hz) ranges (Fig. 3B).

**Figure 3.**
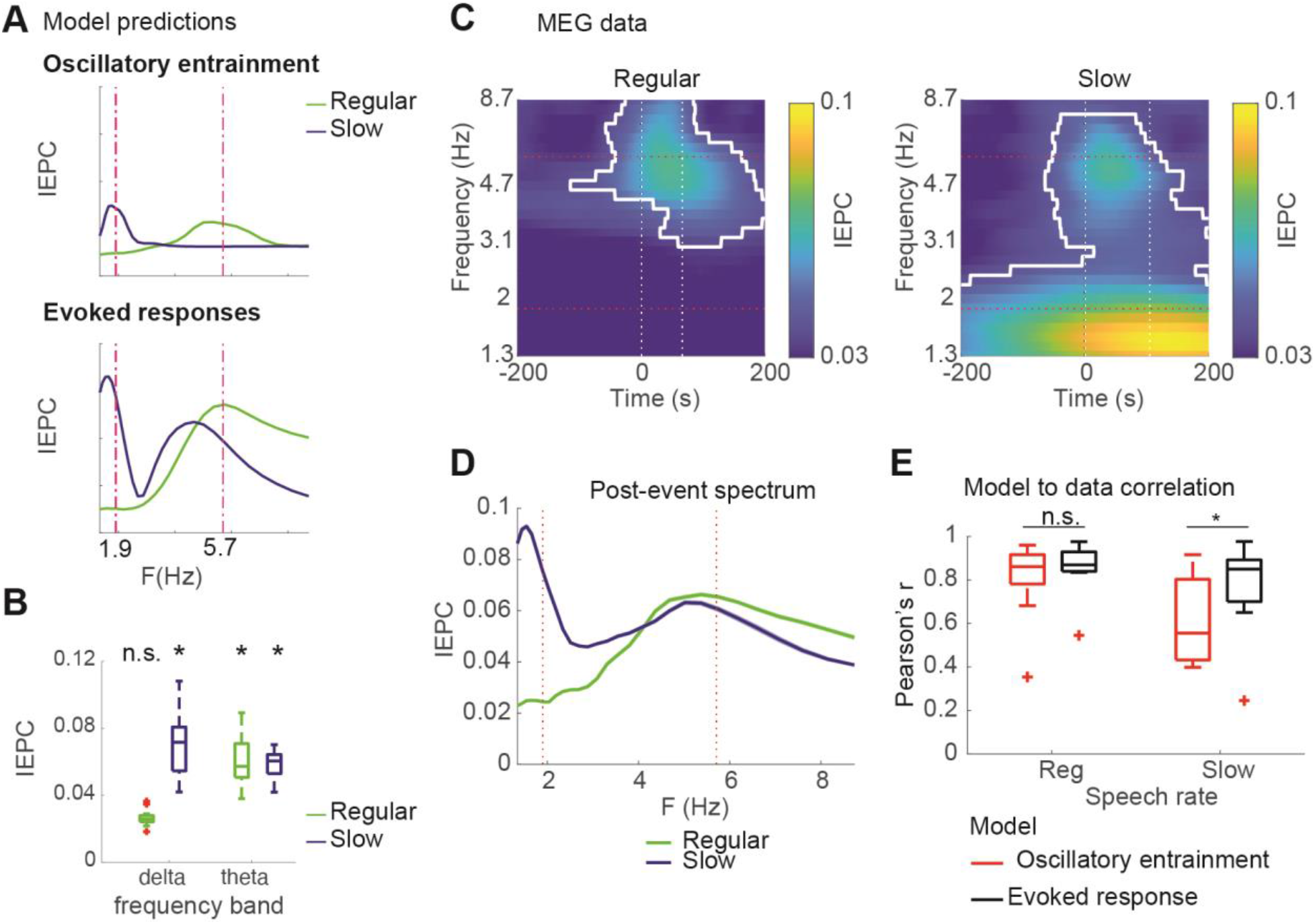
Spectral patterns of IEPC in MEG data. A) Predictions of oscillatory and evoked response models for spectral distribution of phase locking to peakRate events. B) Average IEPC magnitudes observed in regular and slowed speech conditions within time-frequency ROIs in theta and delta bands one oscillatory cycle post peakRate event. C) IEPC patterns observed in MEG responses to speech at regular (left) and slowed (middle) rates. D) Spectral IEPC profile averaged across time corresponds to predictions of the evoked response models (right panel). Significance contours based on 2D cluster-based permutation testing against pre-event baseline, p<.001. E. Correlation between IEPC time courses predicted by the models and observed in the neural data. * *p* < 0.05.

We next turned to testing the two divergent predictions of the two models against MEG data, starting with predictions for spectral distribution. Based on the models’ predictions (Fig. 2 and Fig. 3A), we first took a hypothesis-based approach, testing whether average IEPC values in predefined time-frequency ROIs increased: within a single oscillatory cycle post peakRate event in the theta (4-8 Hz) and delta (1 – 3Hz) ranges (Fig. 3B). In regular speech, we found significant IEPC increase (from theoretical baseline based on Von-Mises distribution) in the theta band (t(11) = 6.9, p < .001, d=2.1), but not the delta band (p > .5), consistent with both models (Fig. 3A). We then turned to the slow speech condition, where the predictions of the two models diverge. We found two spectral peaks in IEPC to peakRate events in slow speech, with a significant increase from baseline in the theta band (t(11) = 8.5, p < .001, d=3.1) and in the delta band (t(11) = 5.2, p < .001, d = 1.9). This pattern is in line with the predictions of the evoked response model but not of the oscillator entrainment model (Fig 3A), as the latter does cannot explain the increased theta IEPC. To verify that these findings did not reflect the specific predefined time-frequency ROIs, we complemented the ROI analysis with a data-driven 2D cluster-based permutation test. This analysis found one cluster in the theta band in the regular speech condition and a large cluster encompassing both theta and delta bands in the slowed speech condition (*p* < 0.001; Fig. 3C, white borders).

Finally, we directly compared how the predictions of both models fit with the spectral IEPC pattern in the data (Fig. 3D for spectral patterns and Fig. 3E for model comparisons). As expected, the difference between models was not significant in the regular speech condition (oscillatory model: mean r = 0.86, evoked response model mean r = 0.81, t(11) = 1.9, p = 0.06). Crucially, in the slowed speech condition, the evoked response model captured the IEPC dynamics significantly better than the oscillatory model (model comparison t (11) = 3.8, p = 0.002), with a large effect size (d = 1.1, post-hoc beta = 0.93). This was because while both models captured the delta-band peak in IEPC, only the evoked response model captured the IEPC dynamics in higher frequencies (oscillatory model: mean r = 0.46, evoked response model mean r = 0.7). Overall, the results of this analysis favor the evoked response model over the oscillatory model.

### Temporal extent of Delta phase locking is limited to a single cycle after peakRate events

We then examined the temporal extent of increased IEPC following peakRate events in the slowed speech condition. The oscillator model predicted that neural IEPC would remain elevated for at least oscillatory cycle, whereas the evoked response model predicted a transient increase in IEPC and return to baseline within 500 ms after the phase reset (Fig. 4A). We calculated IEPC for the MEG data on the same peakRate events as for the model simulations (duration of at least two cycles to subsequent peakRate events), which allowed us to test for continuous entrainment without interference by a subsequent event. We found that IEPC was elevated above baseline for a single cycle following peakRate events, but returned to baseline immediately after (Fig. 4B, cluster-based permutation test against theoretical baseline based on Von-Mises distribution). Notably, this pattern, including the latency of peak IEPC, closely followed the predictions of the evoked response model. Indeed, direct test of the fit of the models’ predictions to the MEG data revealed strong significant correlation with the evoked response model (mean r = 0.59), but not with the oscillator model (mean r = −0.18). This was also reflected in a large significant effect in the direct comparison between models (t(11) = 3.11, p = 0.009, effect size d =0.9, post-hoc power beta = 0.8). Note the negative correlation between data and the oscillatory model, which is due to the reduction in IEPC in the MEG data in the second oscillatory cycle, where IEPC remains high in the oscillatory model. This analysis thus illustrates the transient nature of neural phase locking to peakRate events, which is more consistent with an evoked response mechanism of speech envelope tracking, rather than with an oscillatory entrainment model.

**Figure 4.**
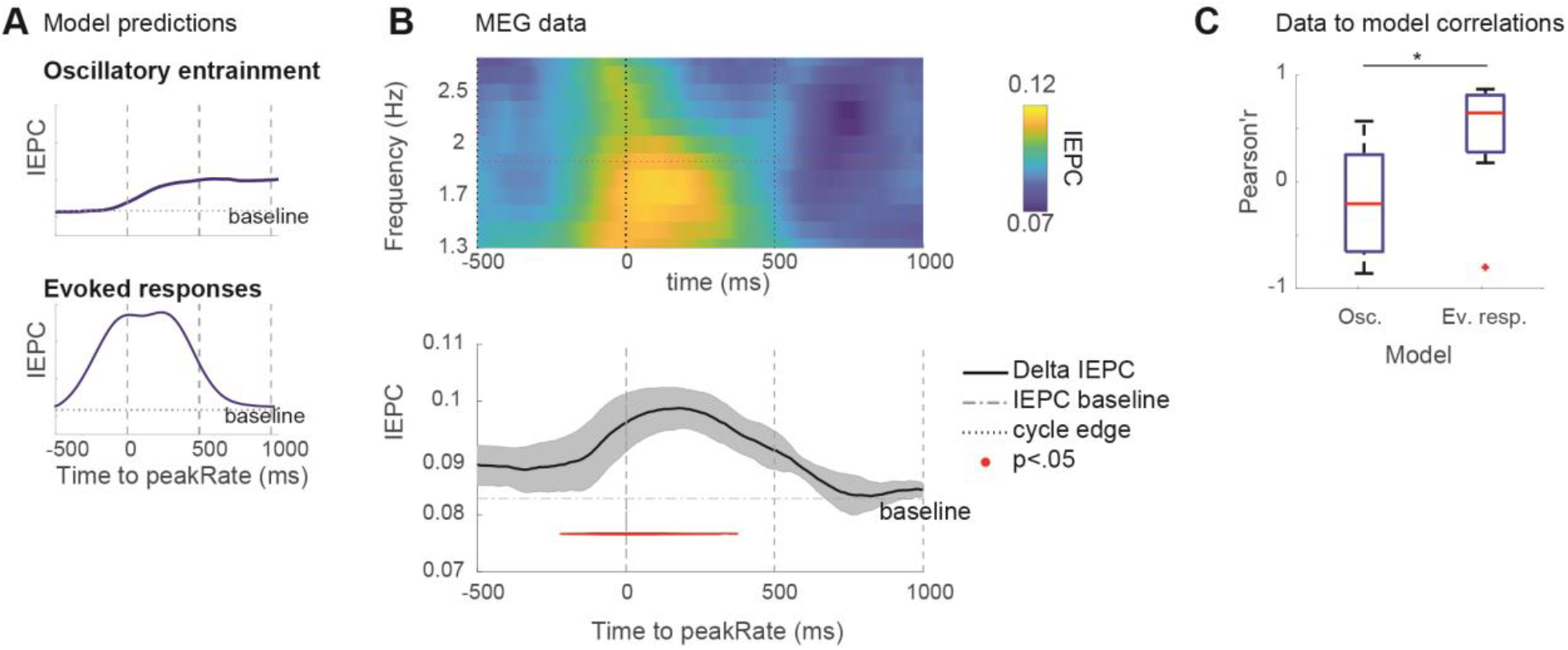
Delta phase locking is limited to a single oscillatory cycle after peakRate events. **A**. Delta IEPC across selected peakRate events that were at least 200 ms away from preceding, and 1000 ms away from subsequent events. White dots mark the extent of one oscillatory cycle in each frequency band. **B**. Delta IEPC time course, red horizontal line marks baseline, red dots mark timepoints of significant deviance from baseline. C. Correlation between IEPC time courses predicted by the models and observed in the neural data. * *p* < 0.05.

Collectively, our findings disagree with an oscillatory entrainment account, which postulates an oscillatory phase-reset after an event, followed by continuous oscillatory reverberation. A more parsimonious account of our results is that the low-frequency phase locking to the speech envelope in MEG is driven by evoked responses to peaks in the envelope rate of change (peakRate). Furthermore, our analysis shows that IEPC to peakRate events reflects the superposition of two different sources: (1) local responses to individual peakRate events and (2) the rate of occurrence of responses to peakRate events. Our analyses also demonstrate that the shift in IEPC frequency bands with changes in speech rate may be the product of a time-frequency decomposition of a series of evoked responses, rather than a shift in the frequency of an entrained oscillator. This finding is a powerful illustration of the importance of explicit computational modeling of alternative neural mechanisms.

In the past, it has been suggested that evoked responses are reduced at slower speech rate, where peakRate magnitudes are smaller, limiting the usability of the evoked response model. In a final analysis we thus tested whether IEPC to peakRate is normalized to account for changes in speech envelope dynamics induced by changes in speech rate.

### Speech rate normalization of peakRate IEPC

The perceptual ability to adapt to variation in the speech signal resulting from changes in the speech rate, i.e., the number of syllables produced per second, is referred to as speech rate normalization. Changes in speech rate results in acoustic changes in the speech signal, including slower amplitude increases at acoustic edges, that is lower peakRate magnitudes (Figure 5A, B). We had previously found that responses to peakRate monotonically scale with peakRate magnitude, being larger for faster changes in the speech amplitude (32). Efficient envelope tracking across speech rates would thus require remapping of neural responses to peakRate magnitude, to account for this overall reduction. Here, we assessed the effect of speech rate on the magnitude of theta IEPC to peakRate events. In the slowed speech, stimuli peakRate magnitudes were 1/3 of those in regular speech (Figures 5C). If no normalization occurs, IEPC magnitudes in slow speech should reflect absolute peakRate values, resulting in an overall reduction in IEPC (Figure 5F, dark dots). In contrast, if theta IEPC to peakRate is invariant to speech rate, it should reflect peakRate values relative to the contextual speech rate, resulting in similar IEPC magnitudes in both speech rate conditions (Figure 5F, light dots).

An evaluation of IEPC after peakRate events, split by peakRate magnitude quantiles, showed comparable theta IEPC in both speech rate conditions (Figure 5D-E), such that average theta IEPC was more robust for larger peakRate magnitudes across both rate conditions (the main effect of peakRate quantile: *b* = 0.01, SD = 0.001, *t* = 1.4, *χ*^2^= 55.0, *p* = 10^−13^). Crucially, they did not differ between regular and slow speech (Interaction effect: *b* = 0.003, SD = 0.005, *t* = 0.6, n.s., Figure 5G), as expected in case of speech rate normalization (Figure 5F, dark dots). Thus, the magnitude of phase reset induced by peakRate depended on its magnitude relative to the local speech rate context, allowing for the flexible encoding of peakRate information at different speech rates.

**Figure 5.**
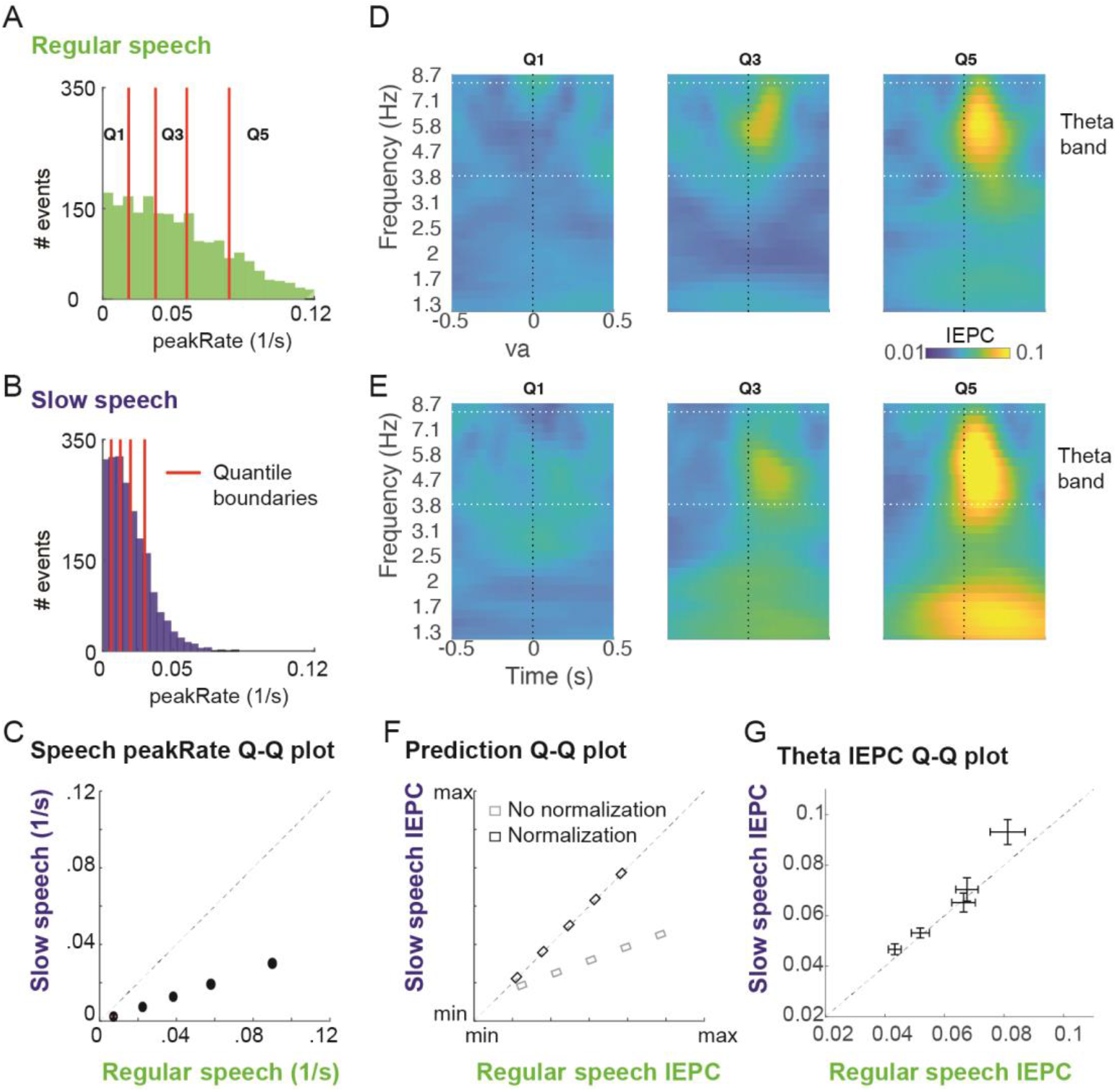
Normalization of peakRate IEPC for contextual speech rate. **A**. Histogram of peakRate magnitudes in regular speech, with quantile boundaries marked in red. **B**. Same as A for slow speech **C**. Quantile-Quantile plot of peakRate magnitudes in regular and slowed speech stimulus. peakRate values in slowed speech stimulus are 1/3 of peakRate values in regular speech stimulus. **D**. IEPC in 1^st^, 3^rd^, 5^th^ peakRate magnitude quantile. Horizontal lines mark the theta frequency range (4-8Hz). **E**. Same as D for slow speech. **F**. Predicted quantile-quantile plots of theta IEPC in regular and slowed speech with (dark) or without (light) normalization. **G**. Quantile-quantile plot of theta-band IEPC (mean, error bars mark ± 1 SEM across subjects) in regular and slow speech. Theta IEPC quantile-quantile values are close to the diagonal, indicating similar magnitudes of theta IEPC in regular and slowed speech conditions.

## Discussion

We evaluated local temporal dynamics in MEG neural representation of the continuous speech envelope under the predictions of oscillatory entrainment and evoked response models, derived from explicit computational models of both processes. Confirming previous results, we found that acoustic edges (peakRate events) drove evoked responses and phase locking over auditory cortical areas (32,38,42). Only the evoked response model captured the spectral and temporal extent of phase-locking to acoustic edges: a transient local component in the theta range, reflective of the evoked response, and – spectrally distinct in slow speech - a separate global component, which captured the frequency of acoustic edges in the stimulus. An analysis of temporally sparse acoustic events further supported the evoked response model: phase locking was transient and limited to the duration of the evoked response. There was no evidence for sustained oscillatory phase locking at the speech rate, as predicted by entrainment models (14,40). Moreover, we found that the magnitude of the evoked phase reset to acoustic edges reflected the speech-rate-normalized amplitude slope at the acoustic edge, offering novel evidence for speech rate normalization. Our results support the central role of acoustic edges in speech comprehension and establish them as the basis for the representation of the speech envelope across methodologies. Overall, our findings confer with the predictions of the evoked response model and suggest that neural phase locking induced by evoked responses to acoustic edges is the primary source of speech envelope tracking in the theta-delta band.

Neural phase resetting may be due to either the superposition of evoked responses or the entrainment of endogenous oscillatory activity. To distinguish between them, we derived the spectral and temporal patterns of phase locking to acoustic edges using simulations of both mechanisms. Model predictions diverged in the slowed speech condition: Spectrally, the evoked response model predicted two spectral peaks in phase reset, in both theta and delta ranges, whereas oscillatory models predicted delta phase locking only. Temporally, the evoked response model predicted only transient phase locking at the speech rate, whereas oscillatory entrainment predicted reverberation: a persisting oscillation for at least 2 cycles after phase-reset (13,14). Note, that the precise temporal extent of IEPC in the oscillator model depends on the decay parameter. However, the hallmark prediction of oscillatory models is that phase-locking will continue after phase-reset beyond a single oscillatory cycle, which is the minimal temporal extend that allows for the model’s proposed functional benefits. It was thus not necessary to include a decay parameter in our models.

In our data, both spectral and temporal patterns of phase locking favored the evoked response model: two spectral peaks and temporally transient phase locking. Notably, both models generated the low frequency phase-locking component in the slow speech condition, corresponding to the frequency of acoustic edge events. While previous work interpreted this component in favor of oscillatory entrainment, our results show that only its temporal extent distinguishes between the two models (43). Overall, our analyses show that a linear convolution of evoked responses to discrete acoustic edge events in speech is sufficient to account for the pattern of neural phase locking to continuous speech. This finding has major implications for theories of speech perception. For instance, instead of oscillatory resonance, predictive processing of speech could rely on non-oscillatory temporal prediction mechanisms guided by statistical learning (16,44,45).

Speech rate normalization is a central behavioral (46,47) and neural phenomenon in speech perception. Shifting of the entrained oscillatory frequency to match the input speech rate was previously proposed as its neural mechanism (23,48). Here, however, we find that the shift of neural phase locking to lower frequencies with speech slowing is an epiphenomenon of spectral analysis of a series of evoked responses. Instead, the magnitude of phase locking to acoustic edges was normalized relative to the distribution of peakRate magnitudes at each rate. Namely, phase locking was comparable across speech rates, despite flatter acoustic edges in slow speech. This suggests that the cortical representations of acoustic edges reflect the magnitude of an edge relative to the contextual speech rate. Such shifting of the dynamic range for acoustic edge magnitudes constitutes a flexible mechanism that maximizes the sensitivity to speech temporal dynamics (49,50) and might not be limited to speech sounds. At the circuit level, the local distribution of peakRate values might be continuously learned via dynamic adjustment of synaptic weights.

Our approach represents a methodological departure from previous investigations of speech envelope tracking. Namely, previous studies focused on cerebro-acoustic coherence (CAC), which reflects the consistency of phase differences between the neural signal and the acoustic stimulus across time (6). CAC is primarily sensitive to regularities across time, such as the rate of phase resets. In contrast, we used inter-event phase coherence (IEPC), which focuses on assessing temporally local similarities in neural phase across repeated occurrences of the same acoustic event (see (51) for IEPC to speech onsets). Our approach revealed that both local phase resets and their rate of occurrence are reflected in IEPC to acoustic edges. In regular speech, both components overlapped within the theta -band. In contrast, slowing of the speech signal separated them in an evoked response component in the theta band, and a slower event frequency component.

Speech rate manipulations are frequently used to study speech envelope tracking (3,24,25,52). Most previous studies used compressed speech to study temporal boundaries on envelope tracking and intelligibility. In contrast, here we used slowed speech to spread distinct acoustic envelope features out in time. It is essential to reconsider previous findings under the evoked response framework. For example, while envelope tracking and intelligibility deteriorate for speech rates higher than 8 Hz, insertion of brief silence periods in compressed speech, which returns the effective speech rate to below 8 Hz, improves intelligibility (52). While this result is typically interpreted as evidence for oscillatory envelope tracking in the theta range, within an evoked response framework it might be reflective of the minimal refractory period of neural populations that encode acoustic edges in speech.

Recent studies have demonstrated that neural responses to speech in M/EEG reflect not only the amplitude envelope but also phonetic and semantic information (53–55). These studies focused on impulse responses (evoked responses) to speech features and their modulation by higher-order information content. We extend upon these studies by pinpointing the envelope feature that drives neural responses to the envelope and show that neural phase-locking to this feature, acoustic edges, can be fully driven by an impulse response. Furthermore, we found responses to speech onsets and amplitude modulations in an ongoing speech on the same sensors over temporal cortical areas. This stands in contrast to intracranial results, which showed that distinct areas of the superior temporal gyrus represent speech onsets and the content of ongoing speech (32,56). However, this is not surprising as we focus on an analysis of MEG data from sensors over the temporal cortex, reflecting the summation of neural sources across auditory cortical areas, such that the shape of the evoked responses to acoustic edges is likely to also be influenced by other temporally confounded events in the speech envelope (such as vowel onsets).

Natural speech does not have a robust temporal rhythmicity (57). Our focus on envelope tracking for natural speech indicates that in this case, neural signatures of envelope tracking are well explained by an evoked response model without the need for an oscillatory component. These results seemingly contradict recent findings of predictive entrainment to music (58). However, our study employed natural speech with considerable variability in inter-edge intervals, unlike in rhythmic musical stimuli. Critically, recent neuropsychological work dissociated neural mechanisms for prediction based on rhythmic streams from predictions in non-rhythmic streams (59). This adds an important caveat to the current debate, suggesting that previous results may perhaps not extend to natural speech with inherent temporal variability and reduced rhythmicity. The present study thus calls to reevaluate the role of oscillatory entrainment in natural speech comprehension. However, it does not preclude the possibility that the introduction of additional rhythmicity to speech, e.g., in poetry or song, recruits additional neural processes associated with the processing of rhythms. Such additional processes might support speech comprehension and could underlie some of the recent findings obtained with a rhythmic speech stimulus (10,21,60). On the other hand, while intelligibility and phase patterns are affected by increased speech rhythmicity or concurrent rhythmic brain stimulation, such findings indicate that oscillations may enhance speech processing, but not that they are necessary for the representation of the significantly less periodic natural speech. Therefore, caution needs to be exercised when extending findings from rhythmic stimuli (e.g., (10,58,61)) to natural speech.

Overall, our results show that an evoked response model accounts for the main neural signatures of speech envelope tracking in MEG. This neural representation of acoustic edges informs about speech rate via inter-event intervals. Moreover, the speech rate normalization of these responses renders this mechanism flexibly adaptable to changes in speech rate. Thus, evoked responses to acoustic edges track the syllabic rate in speech and provide a flexible framework for temporal analysis and prediction during speech perception.

## Data and code availability

All custom-written analysis code will be publicly available upon publication on github (https://github.com/ChangLabUcsf/MEG-SlowSpeech). Data will be made available upon request from the corresponding authors.

## Author contributions

Y.O and E.F.C conceived the study; Y.O, K.K and S.N. designed the experiments and analyzed the data; A.B, Y.O and S.N developed and implemented model simulations; K.K., C.C., and A.F collected and preprocessed the data; K.K. and Y.O. wrote the manuscript; K.K., Y.O, A.B., E.F.C, and S.N revised the manuscript.

## Supplemental Online Materials

**Figure S1.**
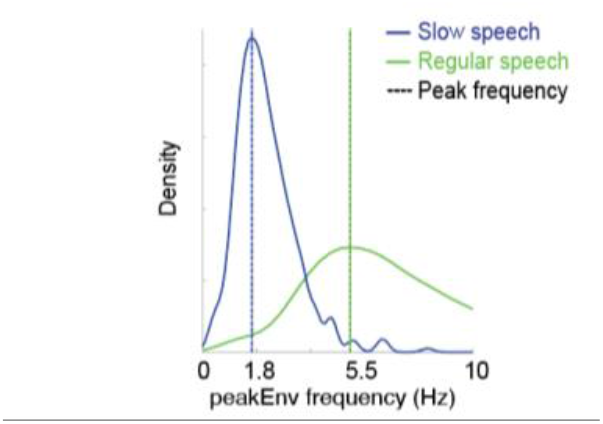
Frequency of peakEnv events. Frequency of peakEnv events in slow speech (blue) and regular speech (green). Dashed vertical lines mark average event frequency in slow (blue, 1.8 Hz) and regular speech (green, 5.5 Hz).

### Evoked low-frequency power following peakRate events

Evoked increase in power is a marker of evoked neural responses and is used to distinguish between evoked responses and oscillatory activity. In addition to calculating the ERP to peakRate events, we thus also tested whether band-passed power would increase after peakRate events. However, we found no significant effects of peakRate on evoked power in theta or delta bands (*p* > 0.05, cluster-based permutation test, data not shown). Our hypothesis that this was due to higher susceptibility of power measures to noise was confirmed in a simulation of the evoked response model (see below).

We hypothesized, that this lack of increase in power in theta or delta bands following peakRate events, this might reflect the high susceptibility of power increases to noise. To assess the effect of noise onto power and phase measures, we tested the evoked response model at noise levels of 1 to 10 relative to response magnitude. We evaluated the effect of noise onto power and IEPC in the theta band (4-8Hz) in the window of a single cycle for a given frequency band after event onset. The effects of noise on power and IEPC were compared using two-sided paired *t*-tests at each noise level (*n* = 20 simulated responses), with Bonferroni correction for the number of comparisons. As predicted, we found continuously large effect sizes for IEPC even at high levels of noise, whereas the effect size for power deteriorated rapidly with the addition of noise.

**Figure S4.**
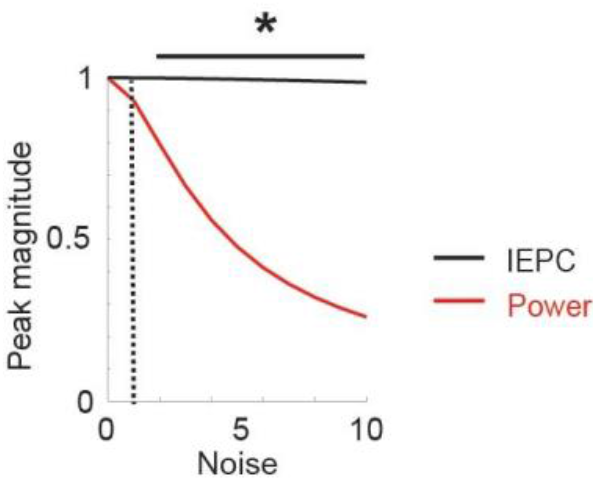
Effect of noise level on IEPC (black) and power (red) after peakRate events in theta band (4-8Hz) for regular speech. * *p* < 0.01

**Figure S5.**
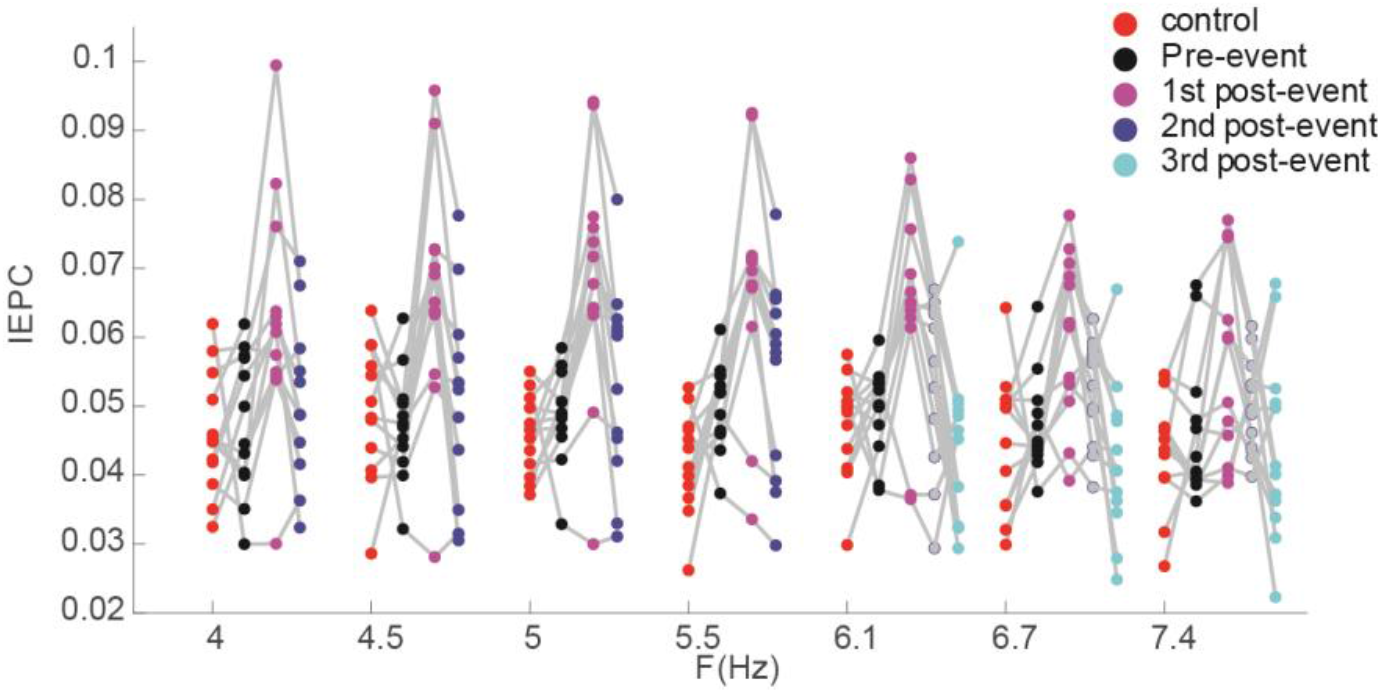
The duration of MEG phase locking is consistent across subjects. Single-subject data for Figure 3B. Average IEPC during baseline (red), pre-event (black), 1st post-event (pink), 2nd post-event (blue), and 3^rd^ post-event cycle (light blue) in each frequency band. IEPC increased from pre-event to 1st post-event and decreased from 1st post-event to 2nd post-event across subjects.

**Figure S6.**
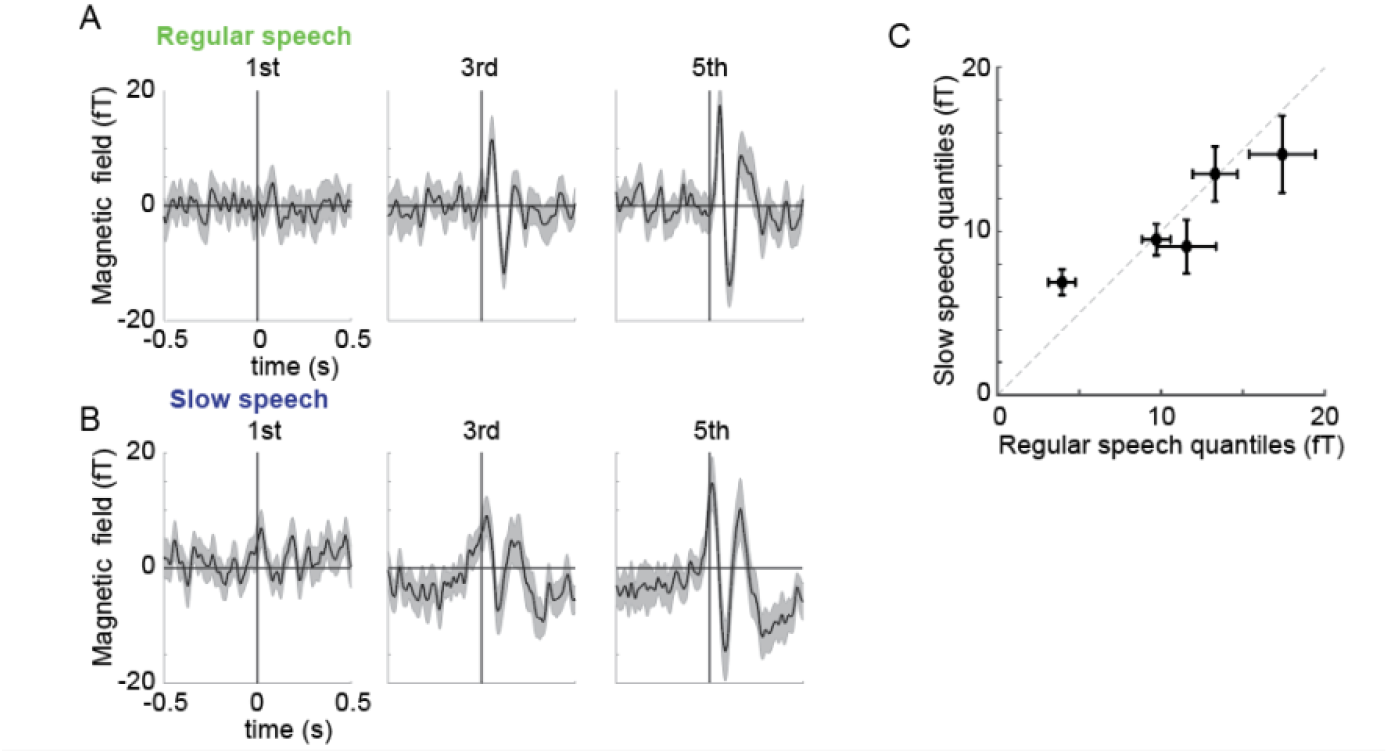
Normalization of peakRate evoked response for contextual speech rate. **A.** Evoked response for peakRate events in 1^st^, 3^rd^, 5^th^ peakRate magnitude quantile. Group-averaged ERPs, error bars are ± 1 SEM. **B**. Same as A for slow speech. **C**. Quantile-quantile plot of peak evoked response (mean (SEM)) in regular and slow speech. Peak evoked response quantile-quantile values are close to the diagonal, indicating the similar distribution of evoked response in regular and slow speech conditions.

**Table S1.** Speech stimulus transcription.

| **Stimulus ID** | **Transcript** |
| --- | --- |
| f1ajrlp1 | Wanted: Chief Justice of the Massachusetts Supreme Court. In April, the S.J.C.'s current leader Edward Hennessy reaches the mandatory retirement age of seventy, and a successor is expected to be named in March. It may be the most important appointment Governor Michael Dukakis makes during the remainder of his administration and one of the toughest. As WBUR's Margo Melnicove reports, Hennessy will be a hard act to follow. |
| f1ajrlp2 | In nineteen seventy-six, Democratic Governor Michael Dukakis fulfilled a campaign promise to de-politicize judicial appointments. He named Republican Edward Hennessy to head the State Supreme Judicial Court. For Hennessy, it was another step along a distinguished career that began as a trial lawyer and led to an appointment as associate Supreme Court Justice in nineteen seventy- one. That year Thomas Maffy, now president of the Massachusetts Bar Association, was Hennessy's law clerk. |
| f1ajrlp3 | The author of more than eight hundred State Supreme Court opinions, Hennessy is widely respected for his legal scholarship and his administrative abilities. Admirers give Hennessy much of the credit for sweeping court reform that began a decade ago, and for last year's legislative approval of thirty-five new judgeships and three hundred million dollars to restore crumbling court houses. Despite the state's massive budget deficit, Hennessy recently urged colleagues in the bar association not to retreat from these hard won gains. |
| f1ajrlp4 | Hennessy is the S.J.C.'s thirty-second chief justice. Holding the court system on the course he has set and plotting it's future agenda won't be an easy job for his successor. |
| f1ajrlp5 | Attorney Haskel Kassler chairs the Judicial Nominating Council, eighteen attorneys and laypeople charged with screening applicants for vacancies on the bench. Usually the J.N.C. refers three nominees to the Governor. His top choice is rated by bar associations and grilled by the Governor's executive council. Kassler says, unlike the Federal Supreme Court, there's no litmus test on particular issues that Massachusetts high court nominees must pass. |
| f1ajrlp6 | All but one of the Chief Justices since eighteen ninety- nine, when Oliver Wendell Holmes was appointed, came from the ranks of S.J.C. associate justices. If he sticks with tradition, Dukakis is likely to elevate one of his appointees to chief. That means Paul Leocos or Ruth Abrams, the only woman on the court. Another possible choice is Herbert Wilkins, a Governor Sargent appointee, and next to Hennessy, the court's most senior member. The other three associate justices were put on the bench by Governor Edward King. And many lawyers say that despite Dukakis' promise to keep the judiciary above the political fray, it's unlikely Dukakis will choose a King appointee to run the state's highest court. For WBUR, I'm Margo Melnicove. |
| m2brrlp1 | Massachusetts may now have the toughest drunken driving law in the nation, thanks to the Safe Roads Act that became law this week. The legislation came complete with a message from the governor to those seeking a little too much comfort and joy this holiday season and beyond. |
| m2brrlp2 | Under the new law the consequences of driving drunk will be swift and severe. Under the old, already tough law, first time offenders used to lose their license for thirty days. Well now they could lose it for as many as ninety. The prison sentence for a second time offender has now doubled. The law is also tougher for vehicular homicide, and it cracks down on minors who drink and drive. State trooper Joseph Holly says this tough new law will mean fewer drunken drivers. |
| m2brrlp3 | Well that's what supporters of the Safe Roads Act are hoping anyway, but will it work? Massachusetts got tough on drunken drivers years ago. What's going to make the difference this time around? Ralph Henksen is chief of the social and behavioral science section at the Boston University school of public health. He says research into what deters the drunken driver shows that often those who drink and drive are risk takers. Even when they're sober they're more likely to run red lights and shun the seat belt. If they think they can drink and drive and get away with it, they'll try. |
| m2brrlp4 | And in fact the state is not planning on putting more police on the road. State officials say there will be the same levels of enforcement. B.U.'s professor Henksen believes the key to the new law's success is its enforcement. |
| m2brrlp5 | The state Safe Roads Act is only a few days old, but it's already facing a challenge. The man who directed the successful effort last election to get the mandatory seat belt law stripped from the law books is now vowing to try to repeal part of this law. Lynn activist Chip Ford has yet to give a name to his committee, but |
|  | he's already set its goal: to repeal what is the most controversial clause in the Safe Roads Act, the per se clause, which allows someone to be punished for drinking and driving before they've even been tried in court. Under the per se part of this new law, if you're arrested, take a breath test, and your blood content level registers point one oh or higher, you can automatically lose your license for ninety days. And that's just at your arraignment. Then you still got to face DWI charges at a trial. State officials say about a quarter of all those arrested for drunken driving are repeat offenders. They say the per se clause will get those people off the road sooner. Chip Ford says it may indeed get them off the road, but it will do it at the expense of their civil rights. |
| m2brrlp6 | Ford and other opponents of the per se clause say it relies exclusively on the breath test, and they say breathalizers are simply unreliable. John Tarantino is a Providence attorney who specializes in drunken driving cases. |
| m2brrlp7 | Tarantino says because of people's physiological differences, the breathalizer test will always be inaccurate on about fourteen percent of the population. Those people will register as having more alcohol in their blood than they actually do. But state officials aren't convinced that's a problem. While they don't say breathalizers are one hundred percent accurate, they do say they are extremely reliable, and if they err at all it's on the conservative side, the side favorable to the driver. As for those who say that taking away someone's license based on the breathalizer results is tantamount to trial by machine, state officials say that simply isn't so. Losing one's license under the per se clause is not a criminal offense. You won't get a criminal record for it. It is a civil infraction and one that they believe will be a key deterrent to the drunken driver. If the per se clause is tested in court, the state says it will pass constitutional muster. And if opponents gather enough signatures to make it a ballot question in nineteen eighty-eight, state officials predict voters will keep it intact. |

**Table S2.**
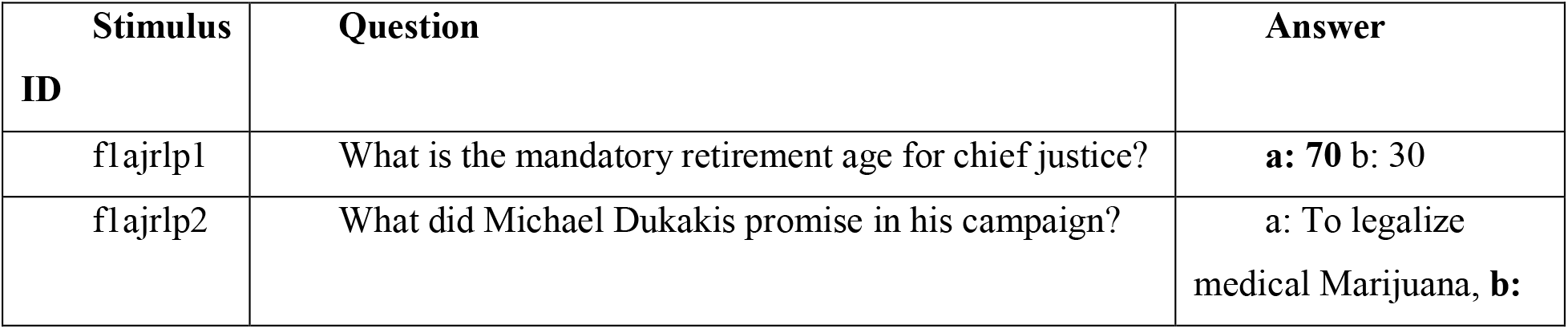
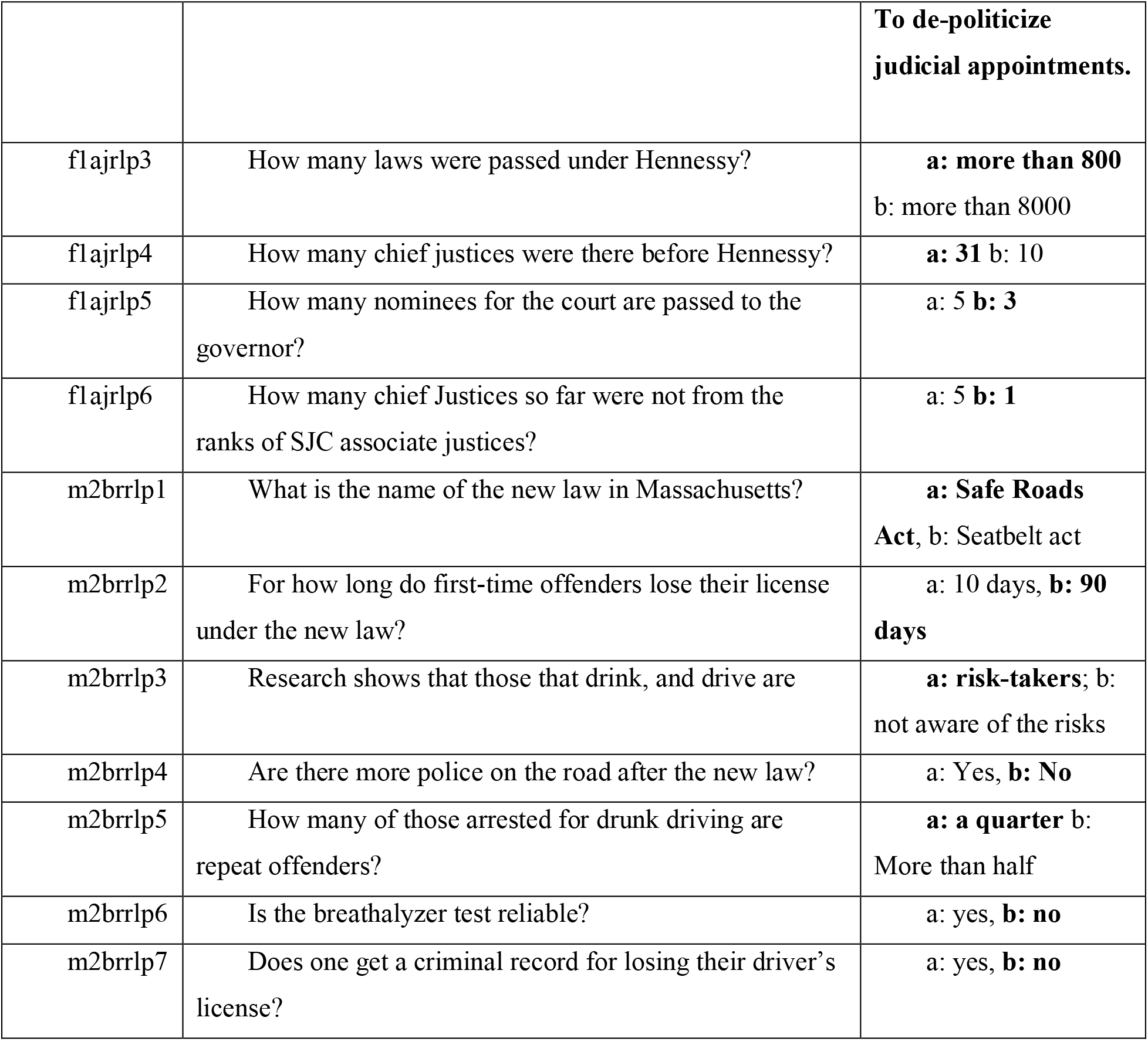
Comprehension questions.

## Notes

### Competing Interest Statement

The authors have declared no competing interest.

### Summary of Updates

Figures 2-4 are revised. The oscillatory model was updated. The temporal extent of delta phase locking was assessed. Assaf Breska was added as a co-author. The authorship order was updated.

